# The effects of combined action observation and motor imagery on corticospinal excitability and movement outcomes: Two meta-analyses

**DOI:** 10.1101/2022.05.23.493106

**Authors:** Samantha Chye, Ashika Chembila Valappil, David J. Wright, Cornelia Frank, David A. Shearer, Christopher J. Tyler, Ceri E. Diss, Omar S. Mian, Neale A. Tillin, Adam M. Bruton

**Affiliations:** Centre for Integrated Research in Life and Health Sciences, School of Life and Health Sciences, University of Roehampton, London, UK, SW15 4JD.; Research Centre for Health, Psychology and Communities, Department of Psychology, Manchester Metropolitan University, Manchester, UK, M15 6GX.; Sports and Movement Research Group, Department of Sports and Movement Science, School of Educational and Cultural Studies, Osnabrück University, Osnabrück, Germany; Faculty of Life Science and Education, University of South Wales, UK, CF37 1DL

**Keywords:** *AOMI*, *motor evoked potentials*, *dual action simulation*, *motor execution*, *motor imagery during action observation*, *transcranial magnetic stimulation*

## Abstract

Motor simulation interventions involving motor imagery (MI) and action observation (AO) have received considerable interest in the behavioral sciences. A growing body of research has focused on using AO and MI simultaneously, termed ‘combined action observation and motor imagery’ (AOMI). The current paper includes two meta-analyses that quantify changes in corticospinal excitability and motor skill performance for AOMI compared to AO, MI and control conditions. Specifically, the first meta-analysis collated and synthesized existing motor evoked potential (MEP) amplitude data from transcranial magnetic stimulation studies and the second meta-analysis collated and synthesized existing movement outcome data from behavioral studies. AOMI had a positive effect compared to control and AO but not MI conditions for both MEP amplitudes and movement outcomes. No methodological factors moderated the effects of AOMI, indicating a robust effect of AOMI across the two outcome variables. The results of the meta-analyses are discussed in relation to existing literature on motor simulation and skill acquisition, before providing viable directions for future research on this topic.

**Highlights:**

- Motor imagery (MI) and action observation (AO) can be combined (AOMI)
- This paper synthesizes neurophysiological and behavioral evidence for AOMI
- AOMI had increased corticospinal excitability compared to AO and control but not MI
- AOMI led to improved movement outcomes compared to AO and control but not MI
- The reported effects of AOMI were maintained across all moderators

## 1. Introduction

According to motor simulation theory (Jeannerod, 1994, 2001, 2006), it is possible to cognitively rehearse an action both overtly and covertly through action observation (AO) and motor imagery (MI), with this cognitive simulation activating motor regions of the brain in a similar manner to physical execution of the action. AO is a bottom-up process that involves the deliberate and structured observation of human movement (Neuman & Gray, 2013), whereas MI is a top-down process that involves the internal generation of the visual and kinesthetic elements of movement (Macintyre et al., 2013). Literature has reported positive behavioral outcomes for AO- and MI-based practice in sport (e.g., Guillot & Collet, 2008; Ste-Marie et al., 2012) and neurorehabilitation settings (e.g., Buccino, 2014; De Vries & Mulder, 2007). There is preliminary evidence indicating that plastic changes in the primary motor system may underpin these behavioral improvements (e.g., Yoxon and Welsh, 2020). A recent large-scale meta-analysis of functional magnetic resonance imaging (fMRI) data reported a shared network including premotor, rostral parietal, and somatosensory areas of the brain active during AO, MI, and movement execution (Hardwick et al., 2018). Notably, there were differences in the neural regions activated during AO and MI, and the brain activity for these overlapped differently with brain activity for physically performed actions. It is possible that using AO and MI simultaneously, typically labelled ‘combined action observation and motor imagery’ (AOMI), may hold greater neural overlap with physical execution.

AOMI refers to a person watching a video or live demonstration of a movement while simultaneously generating, maintaining, and transforming a time-synchronized kinesthetic representation of the same action (Eaves, Riach et al., 2016; Vogt et al., 2013). AOMI has received growing research interest over the last decade and two hypotheses have been proposed to explain why AOMI may be more effective as a motor skill intervention than independent AO or MI interventions. First, Eaves and colleagues (2012, 2014, 2016) suggested the dual action simulation hypothesis (DASH), which proposes that a person will generate separate motor representations for the observed and imagined actions and maintain these as two parallel sensorimotor streams when they engage in AOMI. If a person is simultaneously observing and imagining the same action, these two motor representations are likely to merge as one sensorimotor stream, producing more widespread activity in the premotor cortex compared to AO or MI alone. This is likely due to AOMI increasing activity in shared brain areas for AO and MI, as well as increasing activity in areas solely recruited during AO (e.g., inferior frontal gyrus, ventral premotor area) and MI (e.g., angular gyrus, dorsal premotor area) of an action (Filimon et al., 2015; Hardwick et al., 2018). Second, Meers et al. (2020) introduced the visual guidance hypothesis (VGH) as an alternative account of how AOMI may influence action. They suggest that MI is prioritized during AOMI, and that the AO component might merely serve as an external visual guide that facilitates more vivid MI generation. In contrast to DASH, this would mean that AO does not activate a separate motor representation during AOMI, but rather strengthens the motor representation resulting from MI. Irrespective of the stance taken, both the DASH and VGH suggest that AOMI has the capacity to influence motor skill execution above and beyond AO or MI in isolation through increased activity in motor regions of the brain.

Current neuroscientific evidence using a range of modalities supports this notion, as cortico-motor activity is increased during AOMI of an action compared to independent AO or MI of that same action (Eaves, Riach et al., 2016). Studies using fMRI report distinct neural signatures for AO, MI, and AOMI whereby the blood-oxygen-level-dependent (BOLD) signal is increased and more widespread in the brain regions involved in movement execution when an individual engages in AOMI (e.g., Nedelko et al., 2012; Taube et al., 2015; Villiger et al., 2013). For example, Taube et al. (2015) found greater activation in the supplementary motor area, basal ganglia and cerebellum during AOMI compared to AO, and greater bilateral activity in the cerebellum and greater activation in the precuneus compared to MI. Studies using electroencephalography (EEG) report that AOMI leads to significantly larger event-related desynchronization in the mu/alpha and beta frequency bands, indicative of increased activity over the primary sensorimotor areas of the brain compared to both AO and MI alone (Berends et al., 2013; Eaves, Behmer et al., 2016). These fMRI and EEG findings have important implications for applied practice, where the use of AOMI may prove beneficial in reinforcing motor (re)learning. The increased neural activity during AOMI has the potential to support repetitive Hebbian modulation of intracortical and subcortical excitatory mechanisms through synaptic plasticity, in a similar manner to physical practice, and thus may be an effective method for behavior change (Holmes & Calmels, 2008).

From a neurophysiological perspective, the first meta-analysis in this paper will focus on studies adopting single-pulse transcranial magnetic stimulation (TMS) during AOMI, as this is the most prevalent neuroscientific modality adopted in the literature to date (study count: n = 19 TMS, n = 14 fMRI, n = 10 EEG). When applied to a muscle representation of the primary motor cortex, TMS produces a twitch response in the corresponding muscles called motor evoked potentials (MEPs). MEPs are measured using surface electromyography (EMG) and provide a marker of corticospinal excitability during the time of stimulation (Rothwell, 1997). This approach has been used extensively when studying the neural mechanisms for motor imagery (see e.g., Grosprêtre et al., 2016) and action observation (see e.g., Naish et al., 2014) as it permits a non-invasive assessment of muscle-specific M1 activity and excitability of the whole cortico-spinal pathway that is specific to the final motor command for the action being simulated. Studies using TMS during AOMI have explored changes in MEP amplitudes across a range of movements, including finger movements (Bruton et al., 2020), basketball free throws (Wright, Wood et al., 2018a), walking (Kaneko et al., 2018), and balance movements (Mouthon et al., 2016). Current literature predominantly shows increased corticospinal excitability during AOMI compared to baseline conditions (e.g., Bruton et al., 2020; Wright et al., 2014, 2016). However, studies comparing AOMI against AO or MI have reported increased (e.g., Mouthon et al., 2016; Wright et al., 2016), as well as no differences (e.g., Castro et al., 2021; Mouthon et al., 2015) in corticospinal excitability during AOMI. Given the prevalence of TMS studies exploring the neurophysiological mechanisms underpinning AOMI engagement, it is now possible to synthesize the available MEP amplitude data to quantify the effects of AOMI on corticospinal excitability compared to AO, MI, and control conditions.

AOMI investigations have explored a range of movement outcomes and types, such as movement time for ball rotations (Kawasaki et al., 2018), upper-limb kinematics for dart throwing (Romano-Smith et al., 2019), force production in Nordic hamstring curls (Scott et al., 2018), and radial error from the hole in golf-putting (Marshall & Wright, 2016). This research has been conducted in sport (e.g., Romano-Smith et al., 2019) and rehabilitation (e.g., Scott et al., 2018) contexts, with neurotypical (e.g., Di Rienzo et al., 2019) and neurodivergent (e.g., Marshall et al., 2020) populations. The existing literature has almost exclusively demonstrated that movement outcomes are improved for different motor skills after AOMI interventions when compared to control conditions (e.g., Marshall et al., 2019; Romano-Smith et al., 2018; Shimada et al., 2019).

However, comparisons with AO and MI interventions are equivocal, with some studies showing greater improvements in movement outcomes for AOMI compared to AO (e.g., Bek et al., 2019) and MI interventions (Scott et al., 2018), and other studies showing no such effects (e.g., Romano-Smith et al., 2019) or greater improvements for MI-related interventions (e.g., Marshall & Wright, 2016). Therefore, it is unclear if AOMI should be recommended as the optimal simulation approach when attempting to improve motor skill performance, highlighting the need to synthesize available movement outcome data for AOMI interventions.

Early reviews on AOMI (Eaves, Riach et al., 2016; Vogt et al., 2013) summarized the behavioral neuroscience literature contrasting AO and MI and drew on early AOMI research to support its use as a motor skill intervention. Since then, research has explored changes in MEP amplitudes and movement outcomes associated with engagement in AOMI. This has led to population-specific reviews outlining how AOMI can be used to address sensorimotor deficits for individuals with Parkinson’s disease (Caligiore et al., 2017), developmental coordination disorder (Scott et al., 2021), or during post-stroke rehabilitation (Emerson et al., 2018). Systematic reviews and meta-analyses have synthesized the respective effects of MI and AO on corticospinal excitability (e.g., Grosprêtre et al., 2016; Naish et al., 2014) and motor skill performance (e.g., Ashford et al., 2006; Simonsmeier et al., 2021, Toth et al., 2020) when used in isolation, however, to our knowledge, no such meta-analyses exist for AOMI interventions.

The current paper included two meta-analyses to quantify changes in corticospinal excitability and motor skill performance for AOMI. The first meta-analysis (referred to as MEP meta-analysis) collated and synthesized existing motor evoked potential (MEP) amplitude data from transcranial magnetic stimulation studies as an indicator of corticospinal excitability during AOMI engagement. The second meta-analysis (referred to as Movement meta-analysis) collated and synthesized existing movement outcome data from behavioral studies to assess changes in motor skill performance that result from AOMI interventions. The primary aim for both meta-analyses was to establish the effectiveness of AOMI by comparing its effects on MEP amplitudes (first meta-analysis) and movement outcomes (second meta-analysis) against those for AO, MI, and control conditions. Based on previous literature (see Eaves, Riach et al., 2016), it was hypothesized that AOMI would have a small positive effect when compared to independent AO or MI, and a medium positive effect when compared to control conditions for both meta-analyses. The secondary aim of this paper was to explore several methodological parameters hypothesized to have a moderating effect on the impact of AOMI interventions on MEP amplitudes or movement outcomes across the two meta-analyses. Accounting for the influence of these methodological aspects directly addresses questions raised in early reviews on AOMI (Eaves, Riach et al., 2016; Vogt et al., 2013) and may permit optimal delivery of AOMI interventions in the future.

## 2. Method

### 2.1. Study Protocol

The study procedures were performed according to the methodological guidelines highlighted by the Preferred Reporting Items for Systematic Reviews and Meta-Analyses (PRISMA) statement (Page et al., 2021). The justification for the two meta-analyses and proposed procedures were documented in a pre-registration protocol document (https://osf.io/c68ju?view_only=e6ab97909a6f4f4c8f4323390b3b3c76) that is stored alongside additional supplementary files (https://osf.io/9yebv) on the Open Science Framework. The approach to the two meta-analyses were similar, and the pre-screening literature search was identical (2.1.1). However, the two meta-analyses differed in several ways, including but not limited to the inclusion criteria, effect size preparation, and consideration of moderator effects. A PRISMA flow chart (Figure 1) outlines the study search and selection process for the two meta-analyses.

**Figure 1.**
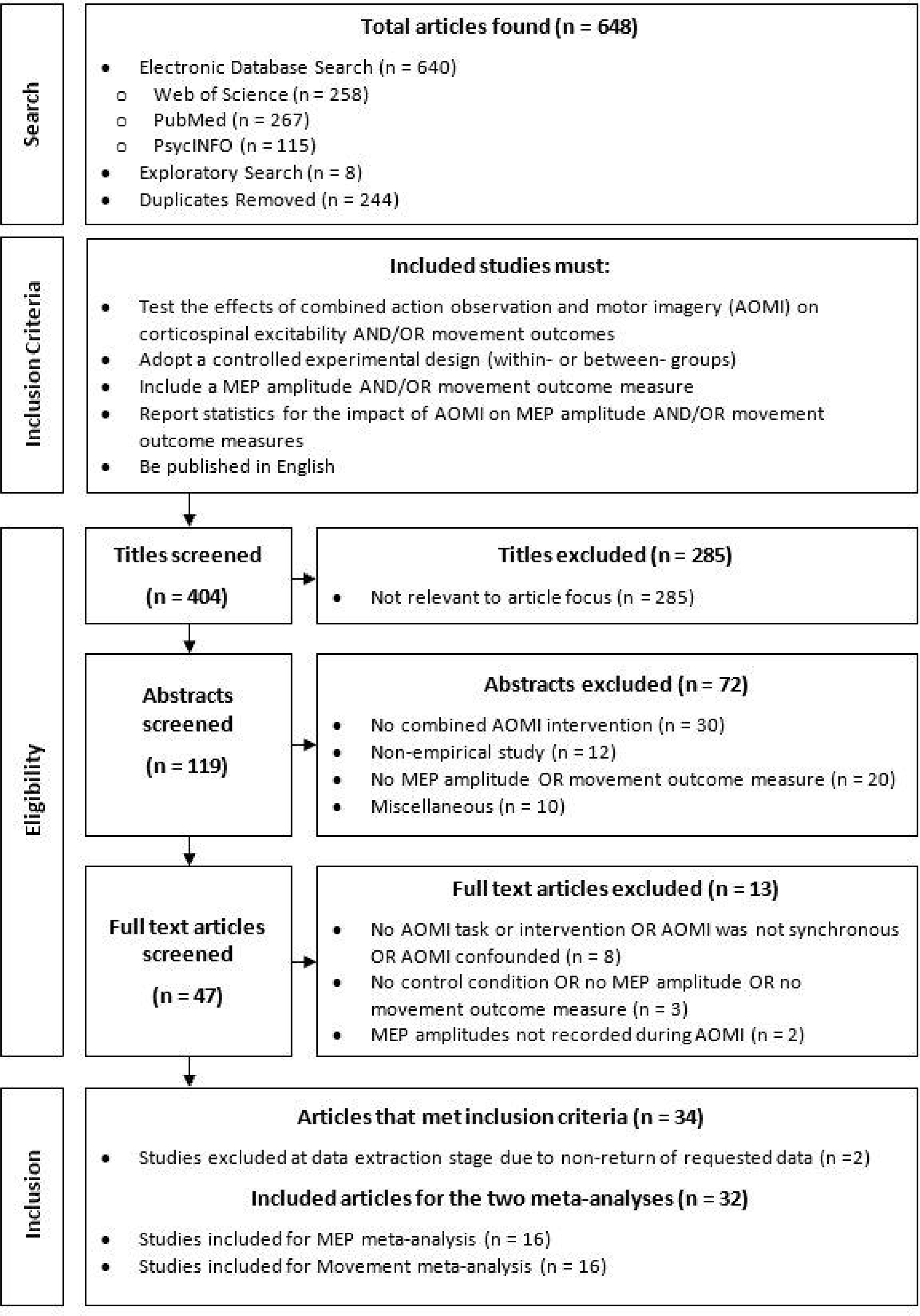
Flow chart of the literature search and selection procedures adopted for the MEP meta-analysis and Movement meta-analysis.

#### 2.1.1. Literature Search and Selection

The literature search was performed using three online databases: PubMed, Web of Science, and PsycINFO. The original search was run in August 2020 and the final search was run in June 2021. Consistent search terms were decided upon and adapted for each database based on the search string requirements (see supplementary files for full search string information). Web of Science was used as an initial search point, followed by PubMed and PsycINFO to cover the biomedical and psychological literature, respectively. The search was limited to studies with human populations that had been published in a peer-reviewed journal in English language, without any limitations to publication years. The literature search with the respective keyword combinations and restrictions provided 396 hits in total, after 244 duplicates were removed. An exploratory search of review articles and prior knowledge of research led to 8 additional papers being added, which took the final number of papers being screened to 404 across the two meta-analyses.

#### 2.1.2. Inclusion of Studies

The decision to include a study was based on five criteria. (1) The study had to either test the effects of AOMI on corticospinal excitability by recording MEP amplitudes during single pulse TMS (MEP meta-analysis) or test the effects of AOMI on movement outcomes by recording parameters related to motor task performance (Movement meta-analysis). Participants were required to engage in an AOMI task as their intervention or experimental method, and AOMI had to be delivered in a synchronous manner (i.e., engagement in both forms of simulation at the same time). Studies with independent AO or MI conditions were included, but only if there was a comparison with an AOMI condition. Studies that included AOMI as an adjunct to physical practice were also included for the Movement meta-analysis, but the inclusion of physical practice was not mandatory, and these studies needed to incorporate physical practice in the control condition to permit assessment of AOMI effects. (2) The study had to adopt a controlled experimental design where the unconfounded effects of the AOMI intervention could be compared within-or between-groups. As such, studies using a correlational or survey design, qualitative research, or case study/single-subject designs were excluded. Studies that did not report a control group or baseline condition were also excluded, unless they contained data from AO or MI conditions that could be compared to the AOMI condition. (3) The study had to include an MEP amplitude (MEP meta-analysis) or movement outcome measure (Movement meta-analysis) with a specific experimental group, or across all study groups. Studies that did not have distinguishable MEP amplitude or movement outcome measures or used neurophysiological modalities other than/combined with TMS (e.g., fMRI, EEG, fNIRS) were excluded unless they also had independent MEP amplitude or movement outcome measures. (4) The study had to report original statistics (i.e., not a re-analysis of already published findings or a review) of the intervention on MEP amplitude or movement outcome measures for the respective meta-analyses. (5) The study had to be published in English.

### 2.2. Screening Process

A three-stage screening process was adopted in this study (i.e., title, abstract, and full-text screening). At all screening stages, the two co-first authors independently screened all articles.

Conflicts were resolved by discussion between the co-first authors in the title screen phase, with additional input from the final author in the abstract and full text screen stages. First, all 404 titles were screened to evaluate relevance to the MEP meta-analysis or Movement meta-analysis, with 119 titles classified as eligible for further inspection of abstracts (intercoder reliability = 94.55%). Second, the 119 abstracts were assessed for eligibility based on the indicative content provided, with 47 abstracts classified as eligible for further inspection as full texts (intercoder reliability = 96.64%). Third, the full text articles were assessed for eligibility based on the study inclusion criteria for the two meta-analyses, with 34 studies progressing to data extraction (intercoder reliability = 97.87%). The mean intercoder agreement across the three stages of the screening process was 96.35%. Whenever data were missing to compute the effect sizes of interest, this information was requested from the corresponding author for the respective article.

### 2.3. Data Processing

#### 2.3.1. Data Extraction

The final 34 studies were split between the two co-first authors based on study focus, with author two (ACV) extracting MEP data from 17 studies for the MEP meta-analysis and author one (SC) extracting movement outcome data from 17 studies for the Movement meta-analysis, respectively. The co-first authors extracted the primary data (i.e., means, standards deviations, and samples sizes) and necessary methodological information to investigate the proposed moderator effects for the present meta-analysis. Once complete, the final author blind-checked the primary data for all studies and contacted the corresponding authors for studies with missing primary data for the two meta-analyses. If the authors did not respond within one month, or prior to the study data analysis cut-off date (January 2021), the article was excluded from the respective meta-analysis. One study was excluded from each of the meta-analyses, leaving 16 studies in the final MEP meta-analysis and 16 studies in the final Movement meta-analysis (see supplementary files for recorded methodological information and primary data for all included articles). No studies met the inclusion criteria for both meta-analyses, meaning the 16 studies synthesized for the MEP meta-analysis were not included in the 16 studies synthesized for the Movement meta-analysis, and vice versa.

#### 2.3.2. Effect Size Preparation

For both meta-analyses, Cohen’s *d* effect sizes were calculated using the mean, standard deviation, and sample size values for the relevant outcome measures prior to conducting the main data analyses using the ‘robu’ function from the ‘robumeta’ package (Fisher et al., 2017) in R studio statistical software (version 2021.09.2 Build 382). The effect size preparation process varied for the MEP meta-analysis and Movement meta-analysis. All studies included in the MEP meta-analysis adopted a repeated measures design, meaning raw data was recorded for all conditions of interest (i.e., AOMI, AO, MI, control). Additional steps were required for studies included in the Movement meta-analysis to account for differences in the movement outcome variables and study designs adopted. When a reduction in a movement outcome measure compared to control was considered an improvement (e.g., reaction time, movement time, mean radial error), the polarity of the calculated effect size was inverted (i.e., positive reversed to negative and vice versa) to ensure an increased value always indicated an improvement.

In order to produce standardized effect sizes that could be compared across the different study designs included in the Movement meta-analysis, effect sizes were calculated for either pre-vs post-test gain comparisons or post-vs post-test comparisons (see supplementary materials for all effect size calculation formulae). Pre-vs-post-test gain comparisons were used to compute effect sizes in studies that adopted mixed experimental designs (i.e., studies that allocated participants to a specific intervention condition and collected data pre- and post-test) as this controls for any pre-test differences that exist between intervention condition groups (Durlak, 2009). Post-vs post-test comparisons were used to compute effect sizes in studies that adopted within-subject (i.e., repeated-measures) or between-subject (i.e., independent intervention condition groups) experimental designs. The number of effect sizes included from each study was not limited as most studies recorded more than one outcome measure for each comparison of interest (i.e., AOMI vs AO, MI, or control). Multiple effect sizes were extracted from studies included in the MEP meta-analysis when MEP amplitudes were collected from more than one target muscle. Multiple effect sizes were extracted from studies included in the Movement meta-analysis when multiple movement outcome measures were recorded. Cohen’s *d* effect size values were converted into Hedge’s *g* effect size values using a small sample size correction formula (Hedges, 1981) for sensitivity analysis.

### 2.4. Data Analysis

Robust variance estimation (RVE) was used to analyze the primary effects for the MEP meta-analysis and Movement meta-analysis. RVE was used because several studies included in each of the two meta-analyses reported multiple relevant effect sizes which were not statistically independent of each other (cf. Tanner-Smith & Tipton, 2014). RVE provides a method for pooling dependent effect size estimates in the absence of any covariance values, mathematically adjusting the standard errors of the effect sizes to account for their dependency (Tanner-Smith & Tipton, 2014; Tanner-Smith et al., 2016). All analyses were run on RStudio using the ‘metafor’ package (Viechtbauer, 2010) and the ‘robumeta’ package (Fisher et al., 2017).

#### 2.4.1. Data Screening

Meta-analyses may be subject to multiple biases (Harrer et al., 2021). To address concerns about publication bias, visual analyses were conducted on the data using funnel plots (Lau et al., 2006), and subsequent statistical analyses were run with the Precision Effect Test (PET) using the RVE approach (Alinaghi & Reed, 2018). For any data that showed publication bias, the trim-and-fill procedure (Duval & Tweedie, 2000) was used to determine the number of unpublished studies required to produce an unbiased estimate of the actual effect size. To address concerns about sample size bias, a sensitivity analysis was run by repeating the main analyses for the two meta-analyses with previously calculated Hedge’s *g* values. This found no meaningful differences in effect size estimate for the primary comparisons, so Cohen’s *d* values are reported in both meta-analyses (cf. Simonsmeier et al., 2021). To address concerns about between-study heterogeneity, outlier diagnostics were completed using the FIND.OUTLIERS function and influence analyses were run using the INFLUENCEANALYSIS function from the ‘dmetar’ package in R (Harrer et al., 2019). This involved visual inspection of “baujat”, “influence” for effect sizes, and “leave-one-out” plots for both effect size and *I^2^* values. Potential outliers and influential effect sizes were identified across the primary comparisons (i.e., AOMI vs AO, AOMI vs MI, AOMI vs Control) for the MEP meta-analysis and Movement meta-analysis, respectively (see supplementary files for detailed overview of data screening results). Removal of the outliers and influential cases had minimal impact on the pooledeffect and heterogeneity estimates for all but one comparison across the two meta-analyses. Two effect sizes were removed from the AOMI vs AO comparison in the Movement meta-analysis (‘delta’ from Frenkel-Toledo et al., 2020; ‘peak angular velocity of the elbow’ from Romano-Smith et al., 2019) as the data points were deemed outliers, influential, and resulted in a meaningful change to the pooled effect and heterogeneity when removed from the RVE data analysis for this comparison. All other effect sizes were retained in the two meta-analyses to preserve the richness of the data.

#### 2.4.2. Quality Assessment

Study quality was assessed to identify if the studies included in the two meta-analyses provide reliably reported data, as well as indicating whether these studies reach an acceptable scientific standard (Borenstein et al., 2021). The first author subjectively assessed the quality of each study using an assessment scale employed in recent meta-analyses (see e.g., Harris et al., 2021). This quality assessment scale was adapted from the Quality Index (Downs & Black, 1998), the Checklist for the Evaluation of Research Articles (Durant, 1994), and the Appraisal Instrument (Genaidy et al., 2007). The quality assessment checklist and individual scores for each study are provided in the supplementary materials.

#### 2.4.3. Primary and Moderator Effects

The primary data analysis for both the MEP meta-analysis and Movement meta-analysis involved correlational RVE models run using the ‘robumeta’ package (Fisher et al., 2017) in RStudio. Overall, eleven moderators were chosen across the two meta-analyses based on previous meta-analyses and reviews focused on the effects of MI and AO on corticospinal excitability (e.g., Grosprêtre et al., 2016; Naish et al., 2014) and motor skill performance (e.g., Ashford et al., 2006; Simonsmeier et al., 2021, Toth et al., 2020). Five moderators were shared across the two meta-analyses (*action observation perspective, skill classification, guided attentional focus, kinesthetic imagery ability, and age*). Three moderators were specific to the MEP meta-analysis (*timing of TMS delivery, number of TMS trials, intensity of TMS pulses*). Five moderators were specific to the Movement meta-analysis (*population type, physical practice, incorporation of PETTLEP principles, context,* and *intervention volume*). Subgroup analyses and meta regressions were used to examine if these moderators influenced the effects of AOMI compared to aggregate data from the AO, MI, and control groups/conditions for the two meta-analyses. Subgroup analyses were used to compare the effects of AOMI based on moderators that consisted of nominal data across the two meta-analyses. The sub-group analyses for the MEP meta-analysis included *action observation perspective, skill classification, guided attentional focus, timing of TMS delivery* moderators*. The sub-group analyses for the Movement meta-analysis included population type, action observation perspective, skill classification, guided attentional focus, physical practice, incorporation of PETTLEP principles, and context* moderators. Meta regression analyses were used to assess if moderators that consisted of interval data predicted the effects of AOMI. Meta regression analyses for the MEP meta-analysis included kinesthetic *imagery ability, age, number of TMS trials, and intensity of TMS pulses.* Meta regression analyses for the Movement meta-analysis included kinesthetic *imagery ability, age, and intervention volume*.

##### 2.4.3.1. Moderators for Both Meta-Analyses

###### 2.4.3.1.1. Action Observation Perspective

Studies have used first-person perspective AO (e.g., Bruton et al., 2020; Romano-Smith et al., 2019) and third-person perspective AO (e.g., Taube et al., 2014; Wright, Wood et al., 2018a) to examine the effects of AOMI on MEP amplitudes and movement outcomes. Sub-group analyses were used to compare the effects of AOMI using first-person vs third-person AO perspectives for both meta-analyses. First-person perspective AO involved the participant viewing the action as if they were performing it (i.e., through their own eyes) and third-person perspective AO involved the participant viewing the action as if another person video recorded them or another person was performing the action (i.e., filmed from a vantage point away from the body). This was determined by checking written text and visual stimuli included in the article.

###### 2.4.3.1.2. Skill Classification

Diverse motor tasks ranging from finger movements (e.g., Meers et al., 2020) to walking (e.g., Kaneko et al., 2018) have been used in previous AOMI literature.

Sub-group analyses were used to compare the effects of AOMI for fine vs gross and continuous vs discrete motor tasks for both meta-analyses. The target movement presented in the AOMI stimuli was classified using a one-dimensional skill classification approach (Spittle, 2021, p.23). Based on this approach, intricate and precise movements using smaller muscle groups (e.g., finger movements) were classed as fine motor tasks; larger muscle movements typically based on fundamental movement patterns (e.g., balance tasks) were classed as gross motor tasks; repetitive movements that have no distinct beginning or end (e.g., walking) were classed as continuous motor tasks; and movements that have an identifiable beginning and end (e.g., putting a golf ball) were classed as discrete motor tasks. Other skill classification comparisons (e.g., open vs closed skills) were not considered in this moderator category due to a lack of coverage in the synthesized literature.

###### 2.4.3.1.4 Guided Attentional Focus

Studies on AO have demonstrated different effects on movement outcomes (e.g., D’Innocenzo et al., 2016) and MEP amplitudes (e.g., Wright, Wood et al., 2018b) when visual attention is directed, or not, to a specific component of the observed action. More recently, Bruton et al. (2020) showed that allocation of visual attention modulates the effects of AOMI on MEP amplitudes for a finger movement task. Sub-group analyses were used to compare the effects of AOMI for guided attentional focus (i.e., use of instructions to direct attention towards a specific aspect of the observed movement) vs unguided attentional focus (i.e., no such instructions) for both meta-analyses. Studies that did not explicitly state if visual attention was manipulated during AOMI were included in the unguided attentional focus sub-group for each meta-analysis.

###### 2.4.3.1.5. Kinesthetic Imagery Ability

Kinesthetic imagery is the imagery modality instructed during AOMI, and the effects of AOMI on movement outcomes reportedly vary as a function of kinesthetic imagery ability (McNeill et al., 2020). Meta regression analyses were used to assess if there was a relationship between kinesthetic imagery ability score and the effects of AOMI for both meta-analyses. Kinesthetic imagery ability data recorded using valid self-report psychometric scales including the Vividness of Movement Imagery Questionnaire -2 (Roberts et al., 2008), Movement Imagery Questionnaire-3 (Williams et al., 2012) and the Movement Imagery Questionnaire-Revised (Hall & Martin, 1997) were included for moderator analyses. Any studies that used non-validated scales such as a visual analogue scale were excluded from the moderator analyses. The imagery ability data was extracted from the studies and standardized by reverse-scoring any measures that adopted an inverse scoring system such that higher numbers meant better imagery ability, before converting all scores to percentage values based on the range of values attainable for each scale.

###### 2.4.3.1.6. Age

Studies have typically recruited adults ranging from early to middle adulthood when assessing MEP amplitudes during AOMI (mean sample age = 27.07 ± 13.48 years) and movement outcomes after AOMI (mean sample age = 30.89 ± 20.24 years). Studies have shown age-related differences in MEP amplitudes during simulation of actions (e.g., Mouthon et al., 2016) and imagery ability is proposed to decline across the lifespan (e.g., Gulyás et al., 2022), suggesting that age may moderate the effects of AOMI on MEP amplitudes and movement outcomes. Meta regression analyses were used to assess if there was a relationship between participant age and the effects of AOMI for both meta-analyses.

##### 2.4.3.2. Moderators for MEP Meta-Analysis

###### 2.4.3.2.1. Timing of Transcranial Magnetic Stimulation Delivery

AO and MI cause phase-specific changes in corticospinal excitability (see e.g., Grosprêtre et al., 2016; Naish et al., 2014 for reviews). Sub-group analysis was used in the MEP meta-analysis to compare the effects of AOMI on MEP amplitudes for TMS delivered at a random point after movement onset against TMS delivered at a targeted point after movement onset (e.g., at the point of maximum movement of the limb). Timing of TMS stimulation delivery refers to the point at which the single pulse is delivered based on the movement displayed in the visual stimuli during AOMI.

###### 2.4.3.2.2. Number of Transcranial Magnetic Stimulation Trials

The number of TMS trials impacts the reliability of the MEP’s evoked during single-pulse TMS (Goldsworthy et al., 2016). Meta regression analysis was used in the MEP meta-analysis to assess if there was a relationship between the number of TMS trials and the effects of AOMI on MEP amplitudes. This was calculated by recording the number of trials where single-pulse TMS was applied to the participant during AOMI for each study.

###### 2.4.3.2.3. Intensity of Transcranial Magnetic Stimulation Pulses

The intensity of TMS pulses impacts the reliability of the MEP’s evoked during single-pulse TMS (Pellegrini et al., 2018). Meta regression analysis was used in the MEP meta-analysis to assess if there was a relationship between intensity of TMS pulses and the effects of AOMI on MEP amplitudes. TMS stimulation intensity refers to the intensity of the TMS stimulator output relative to the resting motor threshold, that is applied to the participant during AOMI.

##### 2.4.3.3. Moderators for Movement Meta-Analysis

###### 2.4.3.3.1. Population Type

AOMI interventions have been shown to benefit movement outcomes in both neurotypical and neurodivergent populations (e.g., Scott et al., 2019). Sub-group analysis was used in the Movement meta-analysis to compare the effects of AOMI on movement outcomes for these two population types. Neurotypical populations included individuals who are not characterized by neurologically atypical patterns, thoughts, behavior, or diagnoses, and neurodivergent populations included individuals whose neurological development and state are considered atypical.

###### 2.4.3.3.2 Physical Practice

Studies have explored the effects of AOMI interventions on movement outcomes with (e.g., Marshall & Wright, 2016) and without (e.g., Taube et al., 2014) physical practice. Sub-group analysis was used in the Movement meta-analysis to compare the effects of AOMI on movement outcomes when used with physical practice vs without physical practice.

###### 2.4.3.3.3. Incorporation of PETTLEP Principles

Some studies have adhered to PETTLEP principles (Holmes & Collins, 2001) when developing and delivering AOMI interventions (e.g., Romano-Smith et al., 2019). Sub-group analysis was used in the Movement meta-analysis to compare the effects of AOMI on movement outcomes with the inclusion of PETTLEP principles vs without inclusion of PETTLEP principles.

###### 2.4.3.3.4. Context

AOMI interventions have been used to target changes in movement outcomes in sport (e.g., Romano-Smith et al., 2018) and rehabilitation (e.g., Marusic et al., 2018) contexts. Sub-group analysis was used in the Movement meta-analysis to compare the effects of AOMI on movement outcomes for sport vs rehabilitation vs other contexts. Studies were classified as sport-or rehabilitation-focused based on the movement being simulated and performed. Studies including movements that did not clearly fall into sports or rehabilitation contexts (e.g., finger movements, ball rotation tasks) were classified as ‘other’.

###### 2.4.3.3.5. Intervention Volume

Studies have delivered AOMI interventions over short- (e.g., Bek et al., 2019) and longer-term (e.g., Shimada et al., 2019) periods when investigating their effects on movement outcomes. Meta regression analysis was used in the Movement meta-analysis to assess the relationship between intervention volume (total minutes) and the effects of AOMI on movement outcomes.

## 3. Results

### 3.1. Study Characteristics

Overall, the two meta-analyses synthesized 111 effect sizes from 32 studies. Of these, the MEP meta-analysis included 54 effect sizes (*n* = 16 studies) and the Movement meta-analysis included 57 effect sizes (*n* = 16 studies). Studies included across the two meta-analyses were published between 2009 and 2021, with a total sample size of 823 participants split across studies included in the MEP meta-analysis (*n* = 234, 77 females, 92 males, 65 undisclosed) and studies included in the Movement meta-analysis (*n* = 589, 281 females, 308 males). The mean age of participants was 27.07 ± 13.48 years and 30.89 ± 20.24 for the two meta-analyses, respectively.

### 3.2. Study Quality

The study quality assessment indicated that all studies included in the two meta-analyses displayed a high degree of rigor. For studies included in the MEP meta-analysis, the quality assessment scores ranged from 18.75-100%, with a mean of 89.58 ± 22.99%. The most poorly addressed items were ‘providing details of a priori sample size determination’ and ‘consistently reporting effect sizes’ with 18.75% and 43.75% of studies satisfying these criteria, respectively (Figure 2a). For studies included in the Movement meta-analysis, the quality assessment scores ranged from 31.25-100%, with a mean of 92.36 ± 17.36%. The most poorly addressed items were ‘providing details of a priori sample size determination’ and ‘applicability of study results to other relevant populations’ with 31.25% and 68.75% of studies satisfying these criteria, respectively (Figure 2b).

**Figure 2.**
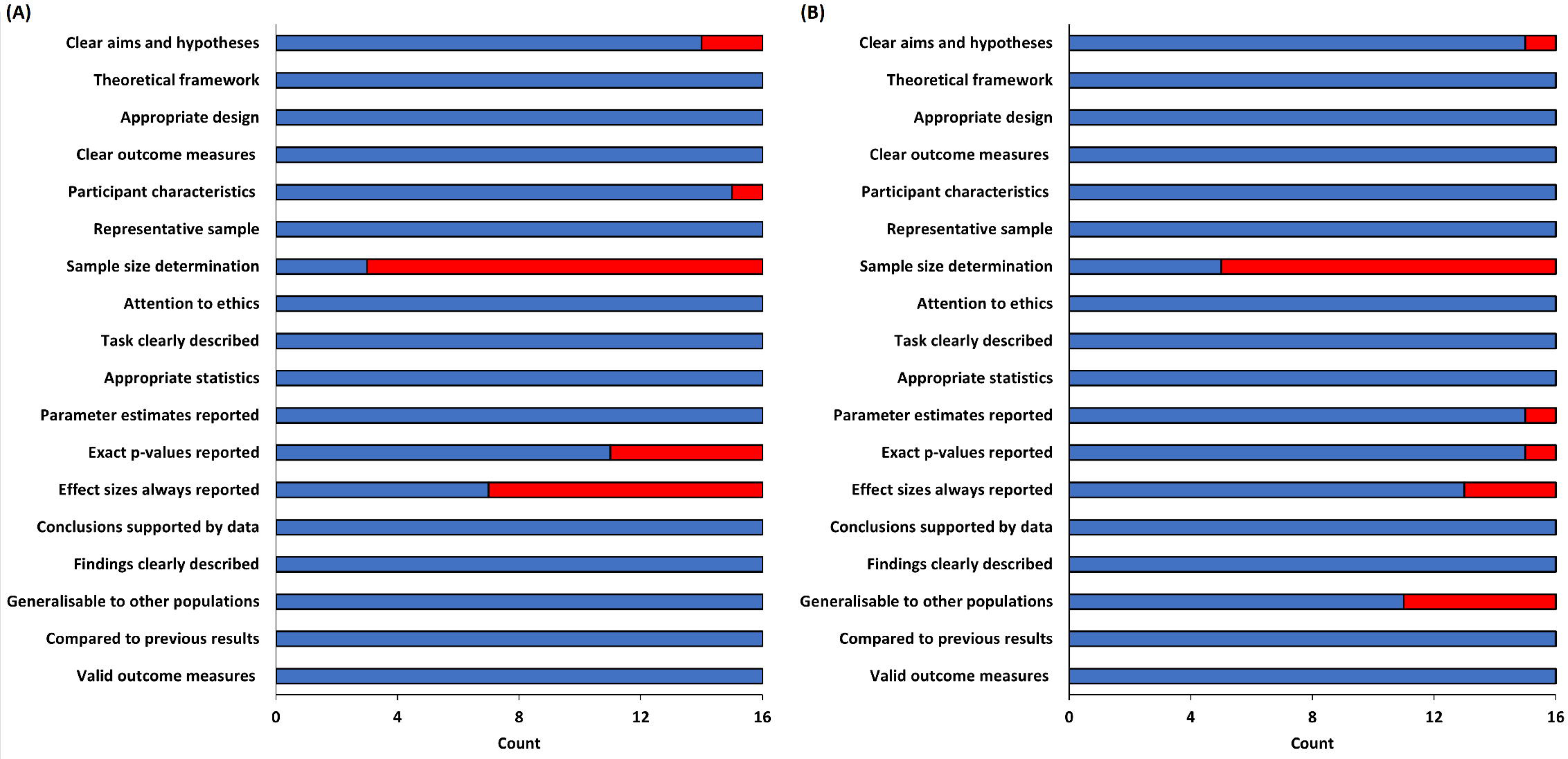
Bar chart displaying quality assessment items and scores for all studies included in the (A) MEP meta-analysis and (B) Movement meta-analysis. The blue bar indicates the number of studies that satisfied, and the red bar indicates the number of studies that did not satisfy, each of the respective quality assessment criteria.

To assess whether studies with smaller samples sizes or lower quality studies were likely to bias the results, meta-regression analyses were run between quality assessment scores and effect size, and sample size and effect size using the ‘*robumeta*’ and ‘metafor’ packages in R. For studies included in the MEP meta-analysis, the analysis used 54 effect sizes from 16 studies and reported that neither quality assessment score (*b* = -0.02, *p* = .25), nor sample size (*b* = -0.01, *p* = .13) predicted the overall effect of AOMI on MEP amplitudes. Similarly, for studies included in the Movement meta-analysis, the analysis used 57 effect sizes from 16 studies and reported that neither quality assessment score (*b* = 0.01, *p* = .76), nor sample size (*b* = -0.03, *p* = .06) predicted the overall effect of AOMI on movement outcomes. The non-significant relationships between quality assessment score, sample size and effect size indicate a low risk of bias for the studies included across the two meta-analyses (*cf*. Harris et al., 2021).

### 3.3. MEP Meta-Analysis

Fifty-four effect sizes from sixteen studies were used in the MEP meta-analysis to determine the overall effect of AOMI on MEP amplitudes. AOMI had a small to medium positive overall effect on MEP amplitudes compared to the control, AO and MI conditions in combination (*d* = 0.48, 95% CI [0.35, 0.61], *p* < .001). The between-study heterogeneity variance was estimated at *τ^2^* = 0.00, with an *I^2^* value of 1.23%. The MEP meta-analysis reported no significant moderators (Tables 1 and 2), demonstrating a robust effect of AOMI on MEP amplitudes irrespective of kinesthetic imagery ability, sample age, intensity of TMS pulses, number of TMS trials, sample age, AO perspective, attentional focus strategy, skill classification, and the timing of TMS delivery.

**Table 1.** Meta-regression analyses for the MEP amplitude data synthesized in the MEP meta- analysis and movement outcome data synthesized in the Movement meta-analysis

| Moderator | N | K | Beta | P | Sig. | $\tau^2$ | $I^2$ |
| --- | --- | --- | --- | --- | --- | --- | --- |
| **MEP Meta-Analysis** |  |  |  |  |  |  |  |
| *Kinesthetic Imagery Ability* | 6 | 19 | -0.01 | **.90** | **No** | 0.00 | 0.00 |
| *Intensity of TMS Pulses* | 15 | 53 | 0.06 | **.13** | **No** | 0.02 | 5.71 |
| *Number of TMS Trials* | 15 | 53 | 0.00 | **.85** | **No** | 0.02 | 11.8 |
| *Age* | 13 | 47 | 0.00 | **.61** | **No** | 0.00 | 0.00 |
| **Movement Meta-Analysis** |  |  |  |  |  |  |  |
| *Intervention Volume* | 14 | 52 | 0.00 | **.72** | **No** | 0.40 | 71.03 |
| *Kinesthetic Imagery Ability* | 5 | 21 | -0.01 | **.86** | **No** | 2.40 | 88.53 |
| *Age* | 16 | 57 | -0.01 | **.19** | **No** | 0.38 | 68.73 |

Note. N = number of studies, K = number of effect sizes, Beta = regression coefficient, P = significance value, Sig. = significance threshold met or not, $\tau^2$ = measure of variation around average, $I^2$ = measure of proportion of observed variance (%).

**Table 2.** Sub-group analyses for the MEP amplitude data synthesized in the MEP meta-analysis

| Sub-group | N | K | SMD | 95% CI |  | P | Sig. | τ<sup>2</sup> | I<sup>2</sup> | Sig. Mod. |
| --- | --- | --- | --- | --- | --- | --- | --- | --- | --- | --- |
| Action Observation Perspective |  |  |  |  |  |  |  |  |  |  |
| First-person | 9 | 23 | 0.50 | 0.33 | 0.67 | <.001 | Yes | 0.00 | 0.00 | ref |
| Third-person | 7 | 31 | 0.48 | 0.12 | 0.84 | .02 | Yes | 0.15 | 45.41 | ns |
| Directed Attentional Focus |  |  |  |  |  |  |  |  |  |  |
| Yes | 1 | 2 | 0.38 | NA | NA | NA | NA | 0.00 | 0.00 | ref |
| No | 15 | 52 | 0.49 | 0.34 | 0.64 | <.001 | Yes | 0.02 | 7.15 | ns |
| Skill Classification 1 |  |  |  |  |  |  |  |  |  |  |
| Fine | 10 | 24 | 0.48 | 0.33 | 0.64 | <.001 | Yes | 0.00 | 0.00 | ns |
| Gross | 6 | 30 | 0.52 | 0.07 | 0.97 | .03 | Yes | 0.19 | 52.98 | ref |
| Skill Classification 2 |  |  |  |  |  |  |  |  |  |  |
| Discrete | 12 | 32 | 0.52 | 0.35 | 0.70 | <.001 | Yes | 0.00 | 0.00 | ns |
| Continuous | 4 | 22 | 0.34 | 0.16 | 0.52 | <.01 | Yes | 0.09 | 33.19 | ref |
| Timing of TMS Delivery |  |  |  |  |  |  |  |  |  |  |
| Targeted | 12 | 35 | 0.47 | 0.30 | 0.64 | <.001 | Yes | 0.00 | 0.00 | ref |
| Random | 4 | 19 | 0.53 | 0.28 | 0.77 | <.001 | Yes | 0.08 | 28.61 | ns |

*Note.* N = number of studies, K = number of effect sizes, SMD = standardized mean difference, 95% CI = lower and upper confidence intervals, P = significance value, Sig. = significance threshold met or not, $\tau^2$ = measure of variation around average, $I^2$ = measure of proportion of observed variance (%), Sig. Mod. = comparison between sub-group categories. NA = insufficient number of data points for the analysis, ref = reference category, ns = no significant difference compared to reference category

#### 3.3.1. AOMI vs Control

Nineteen effect sizes from thirteen studies were used to compare AOMI vs control conditions in the MEP meta-analysis. AOMI had a medium positive effect on MEP amplitudes compared to control conditions (*d* = 0.54, 95% CI [0.41, 0.66], *p* < .001). The between-study heterogeneity variance was estimated at *τ^2^* = 0.00, with an *I^2^* value of 0.00%.

#### 3.3.2. AOMI vs AO

Twenty-three effect sizes from thirteen studies were used to compare AOMI vs AO conditions in the MEP meta-analysis. AOMI had a small to medium positive effect on MEP amplitudes compared to AO conditions (*d* = 0.45, 95% CI [0.27, 0.63], *p* < .001). The between-study heterogeneity variance was estimated at *τ^2^* 0.02, with an *I^2^* value of 10.32%.

#### 3.3.3. AOMI vs MI

Twelve effect sizes from six studies were used to compare AOMI vs MI conditions in the MEP meta-analysis. AOMI had no significant effect on MEP amplitudes compared to MI conditions (*d* = 0.25, 95% CI [-0.13, 0.63], *p* = .14). The between-study heterogeneity variance was estimated at *τ^2^*= 0.11, with an *I^2^* value of 42.92%.

### 3.4. Movement Meta-Analysis

Fifty-seven effect sizes from sixteen studies were used in the Movement meta-analysis to determine the overall effect of AOMI on movement outcomes. AOMI had a small to medium positive overall effect on movement outcomes compared to the control, AO and MI conditions in combination (*d* = 0.48, 95% CI [0.18, 0.78], *p* < .01). The between-study heterogeneity variance was estimated at *τ^2^* = 0.39, with an *I^2^* value of 69.68%. The Movement meta-analysis reported no significant moderators (Tables 1 and 3), demonstrating a robust effect of AOMI on movement outcomes irrespective of intervention volume, kinesthetic imagery ability, sample age, AO perspective, study context, attentional focus strategy, incorporation of PETTLEP, physical practice, population type, and skill classification.

**Table 3.**
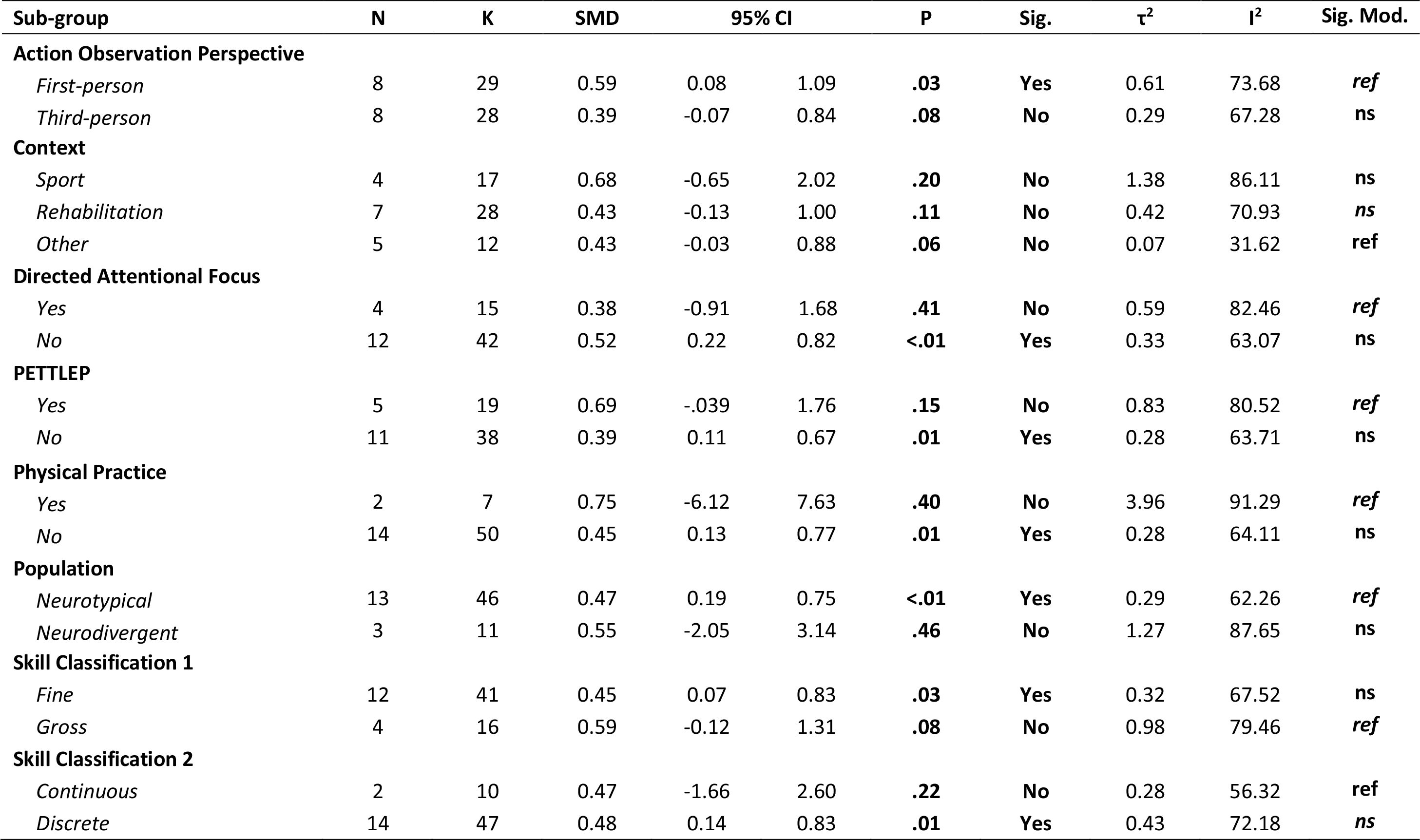
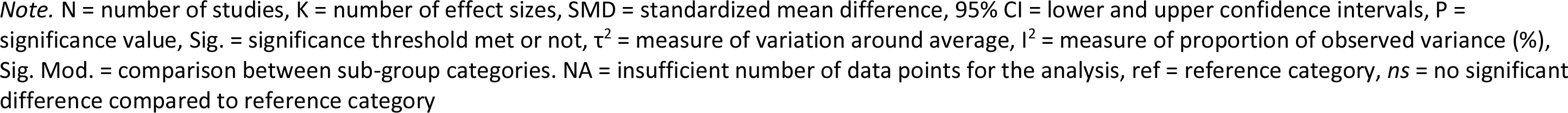
Sub-group analyses for the movement outcome data synthesized in the Movement meta-analysis

| Sub-group | N | K | SMD | 95% CI |  | P | Sig. | τ² | I² | Sig. Mod. |
| --- | --- | --- | --- | --- | --- | --- | --- | --- | --- | --- |
| Action Observation Perspective |  |  |  |  |  |  |  |  |  |  |
| First-person | 8 | 29 | 0.59 | 0.08 | 1.09 | .03 | Yes | 0.61 | 73.68 | ref |
| Third-person | 8 | 28 | 0.39 | -0.07 | 0.84 | .08 | No | 0.29 | 67.28 | ns |
| Context |  |  |  |  |  |  |  |  |  |  |
| Sport | 4 | 17 | 0.68 | -0.65 | 2.02 | .20 | No | 1.38 | 86.11 | ns |
| Rehabilitation | 7 | 28 | 0.43 | -0.13 | 1.00 | .11 | No | 0.42 | 70.93 | ns |
| Other | 5 | 12 | 0.43 | -0.03 | 0.88 | .06 | No | 0.07 | 31.62 | ref |
| Directed Attentional Focus |  |  |  |  |  |  |  |  |  |  |
| Yes | 4 | 15 | 0.38 | -0.91 | 1.68 | .41 | No | 0.59 | 82.46 | ref |
| No | 12 | 42 | 0.52 | 0.22 | 0.82 | <.01 | Yes | 0.33 | 63.07 | ns |
| PETTLEP |  |  |  |  |  |  |  |  |  |  |
| Yes | 5 | 19 | 0.69 | -.039 | 1.76 | .15 | No | 0.83 | 80.52 | ref |
| No | 11 | 38 | 0.39 | 0.11 | 0.67 | .01 | Yes | 0.28 | 63.71 | ns |
| Physical Practice |  |  |  |  |  |  |  |  |  |  |
| Yes | 2 | 7 | 0.75 | -6.12 | 7.63 | .40 | No | 3.96 | 91.29 | ref |
| No | 14 | 50 | 0.45 | 0.13 | 0.77 | .01 | Yes | 0.28 | 64.11 | ns |
| Population |  |  |  |  |  |  |  |  |  |  |
| Neurotypical | 13 | 46 | 0.47 | 0.19 | 0.75 | <.01 | Yes | 0.29 | 62.26 | ref |
| Neurodivergent | 3 | 11 | 0.55 | -2.05 | 3.14 | .46 | No | 1.27 | 87.65 | ns |
| Skill Classification 1 |  |  |  |  |  |  |  |  |  |  |
| Fine | 12 | 41 | 0.45 | 0.07 | 0.83 | .03 | Yes | 0.32 | 67.52 | ns |
| Gross | 4 | 16 | 0.59 | -0.12 | 1.31 | .08 | No | 0.98 | 79.46 | ref |
| Skill Classification 2 |  |  |  |  |  |  |  |  |  |  |
| Continuous | 2 | 10 | 0.47 | -1.66 | 2.60 | .22 | No | 0.28 | 56.32 | ref |
| Discrete | 14 | 47 | 0.48 | 0.14 | 0.83 | .01 | Yes | 0.43 | 72.18 | ns |

*Note.* N = number of studies, K = number of effect sizes, SMD = standardized mean difference, 95% CI = lower and upper confidence intervals, P = significance value, Sig. = significance threshold met or not, $\tau^2$ = measure of variation around average, $I^2$ = measure of proportion of observed variance (%), Sig. Mod. = comparison between sub-group categories. NA = insufficient number of data points for the analysis, ref = reference category, *ns* = no significant difference compared to reference category

#### 3.4.1. AOMI vs Control

Twenty-seven effect sizes from twelve studies were used to compare AOMI vs control conditions in the Movement meta-analysis. AOMI had a medium to large positive effect on movement outcomes compared to control conditions (*d* = 0.67, 95% CI [0.16, 1.18], *p =* .02). The between-study heterogeneity variance was estimated at *τ^2^* = 0.48, with an *I^2^* value of 70.74%.

#### 3.4.2. AOMI vs AO

Nineteen effect sizes from nine studies were used to compare AOMI vs AO conditions in the Movement meta-analysis. AOMI had a small to medium positive effect on movement outcomes compared to AO conditions (d = 0.44, 95% CI [0.07, 0.81], *p* = .03). The between-study heterogeneity variance was estimated at *τ^2^*= 0.24, with an *I^2^* value of 63.23%.

##### 3.3.2.3. AOMI vs MI

Eleven effect sizes from six studies were used to compare AOMI vs MI conditions in the Movement meta-analysis. AOMI had no significant effect on movement outcomes compared to MI conditions (*d* = 0.53, 95% CI [-0.59, 1.66], *p* = .28). The between-study heterogeneity variance was estimated at *τ^2^* = 1.30, with an *I^2^* value of 84.46%.

### 3.5. Publication Bias

Based on the funnel plots of effect size level (see supplementary files), publication bias was identified to be unlikely for all comparisons across the two meta-analyses. Regardless, we ran trim and fill and PET analyses using the RVE method to retrieve an unbiased effect size estimate corrected for publication bias for all comparisons made in the MEP meta-analysis and Movement meta-analysis.

For the control comparison in the MEP meta-analysis, both the PET-intercept (*b0* = 0.09, *p* = 0.83) and the PET-slope (*b1* = 1.21, *p* = 0.32) were not statistically significant, suggesting that publication bias was unlikely. For the AO comparison in the MEP meta-analysis, the funnel plot of effect size level (Figure 9, left) indicated asymmetry. Trim-and-fill analysis proposed 4 missing values (Figure 9, right), and the effect size changed from a medium (*d* = 0.54) to a small to medium positive effect (*d* = 0.43), suggesting minimal effects of publication bias for this dataset. For the PET, both the PET-intercept (*b0* = -1.00, *p* = 0.16) and the PET-slope (*b1* = 3.85, *p* = 0.07) were not statistically significant, suggesting that publication bias was unlikely. For the MI comparison in the MEP meta-analysis, the funnel plots indicated no funnel asymmetry and the trim-and-fill analysis reported zero missing values. For the PET, both the PET-intercept (*b0* = -1.14, *p* = 0.43) and the PET-slope (*b1* = 3.59, *p* = 0.41) were not statistically significant, suggesting that publication bias was unlikely.

**Figure 3.**
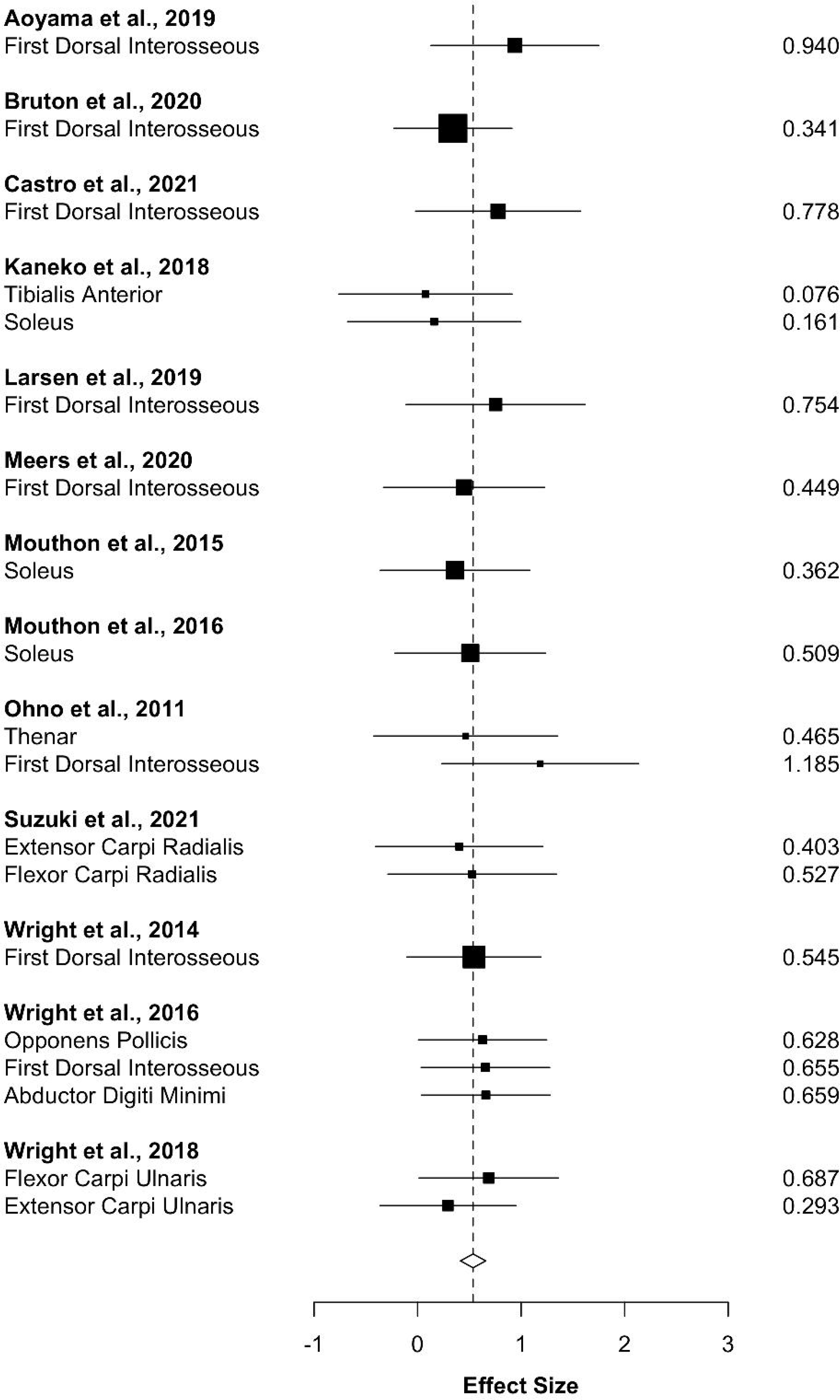
Forest plot of effect sizes (*d*) for all studies included in the AOMI vs control condition comparison for the MEP meta-analysis. The combined estimate (dashed vertical line) and 95% confidence interval (hollow diamond) indicates AOMI has a medium positive effect on MEP amplitudes compared to control conditions. The size of each black square indicates the weight of the study effect size in the combined analysis. Multiple effect sizes are reported for a study if it recorded MEP amplitude data from more than one target muscle.

**Figure 4.**
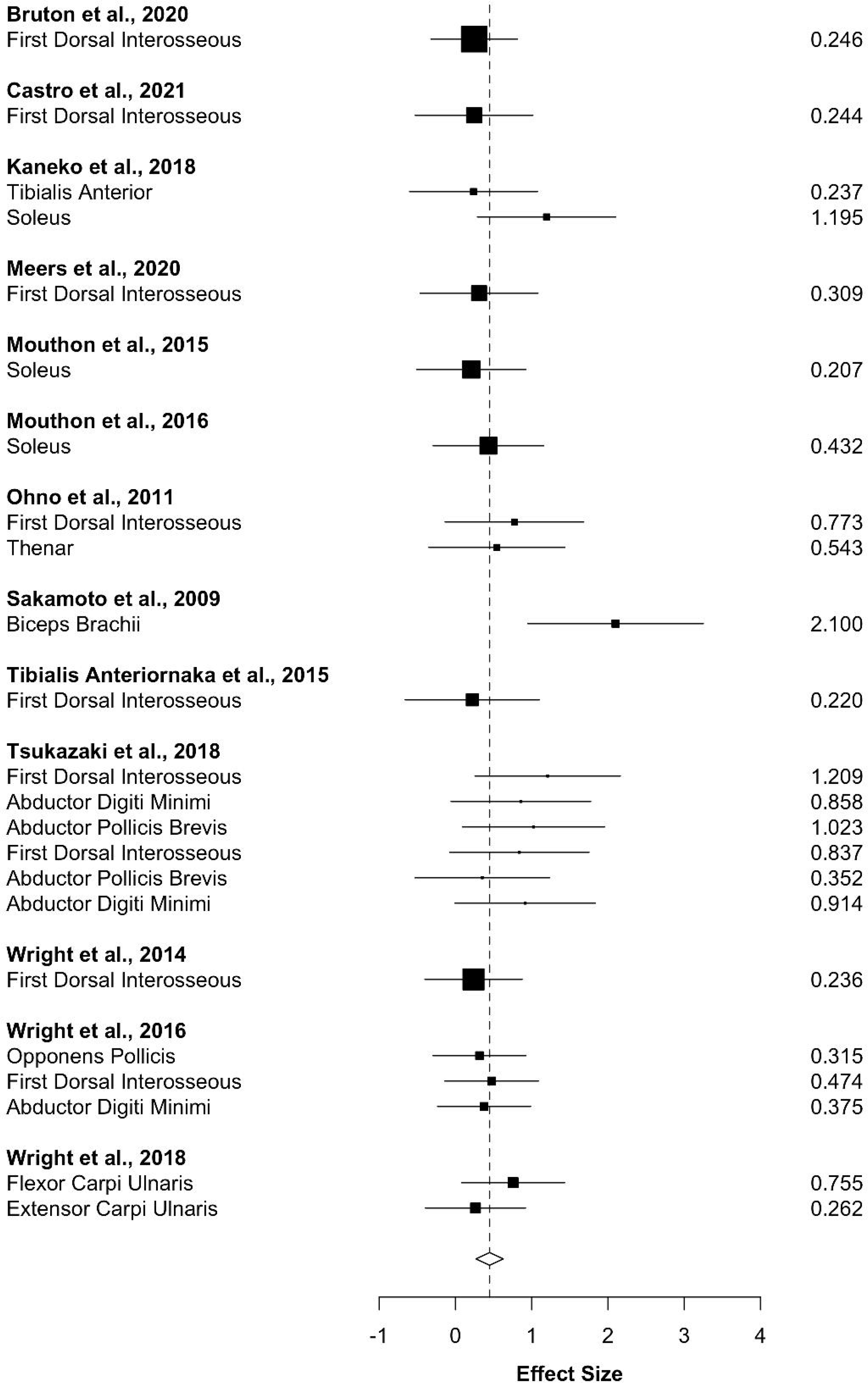
Forest plot of effect sizes (*d*) for all studies included in the AOMI vs AO condition comparison for the MEP meta-analysis. The combined estimate (dashed vertical line) and 95% confidence interval (hollow diamond) indicates AOMI has a small to medium positive effect on MEP amplitudes compared to AO conditions. The size of each black square indicates the weight of the study effect size in the combined analysis. Multiple effect sizes are reported for a study if it recorded MEP amplitude data from more than one target muscle.

**Figure 5.**
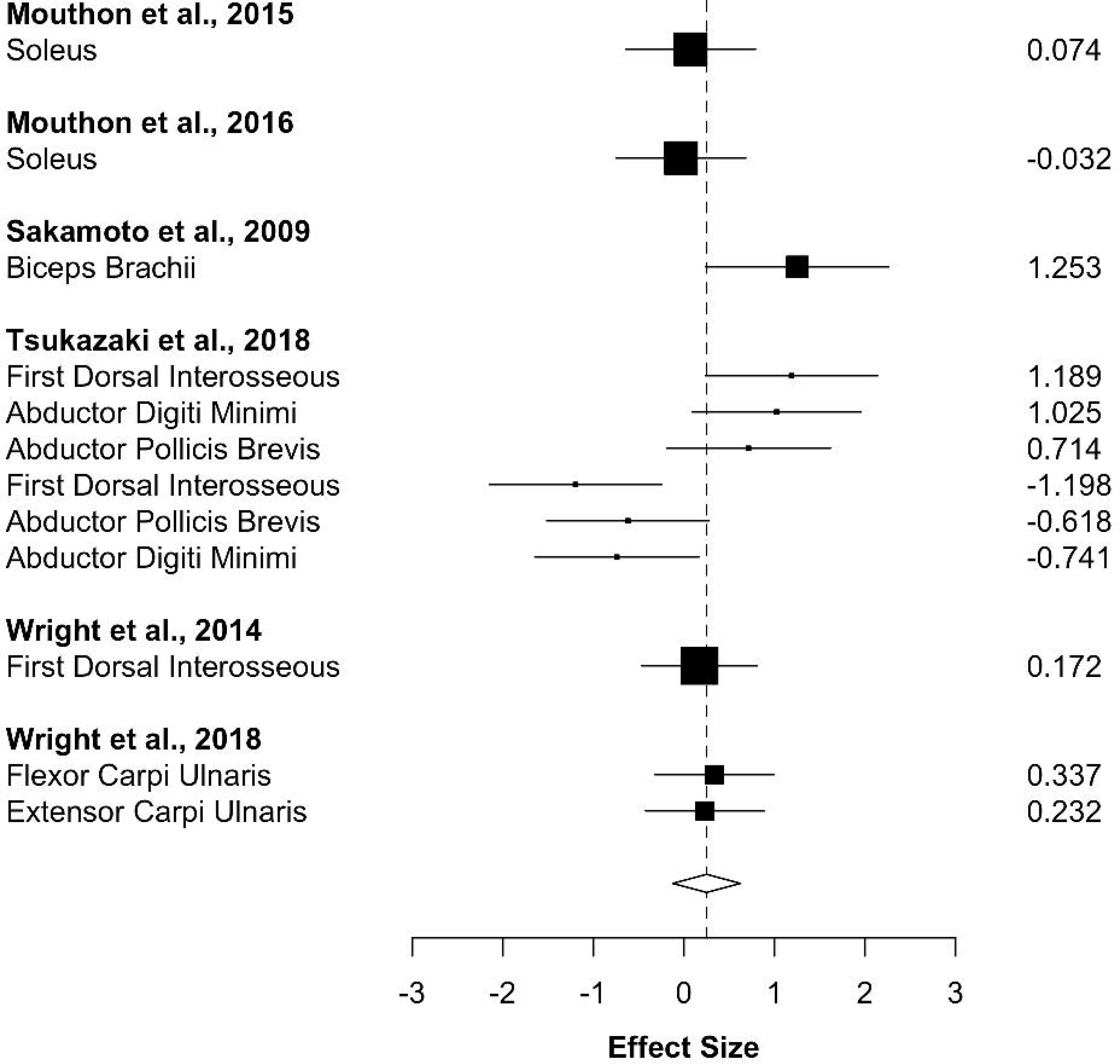
Forest plot of effect sizes (*d*) for all studies included in the AOMI vs MI condition comparison for the MEP meta-analysis. The combined estimate (dashed vertical line) and 95% confidence interval (hollow diamond) indicates AOMI has no significant effect on MEP amplitudes compared to MI conditions. The size of each black square indicates the weight of the study effect size in the combined analysis. Multiple effect sizes are reported for a study if it recorded MEP amplitude data from more than one target muscle.

**Figure 6.**
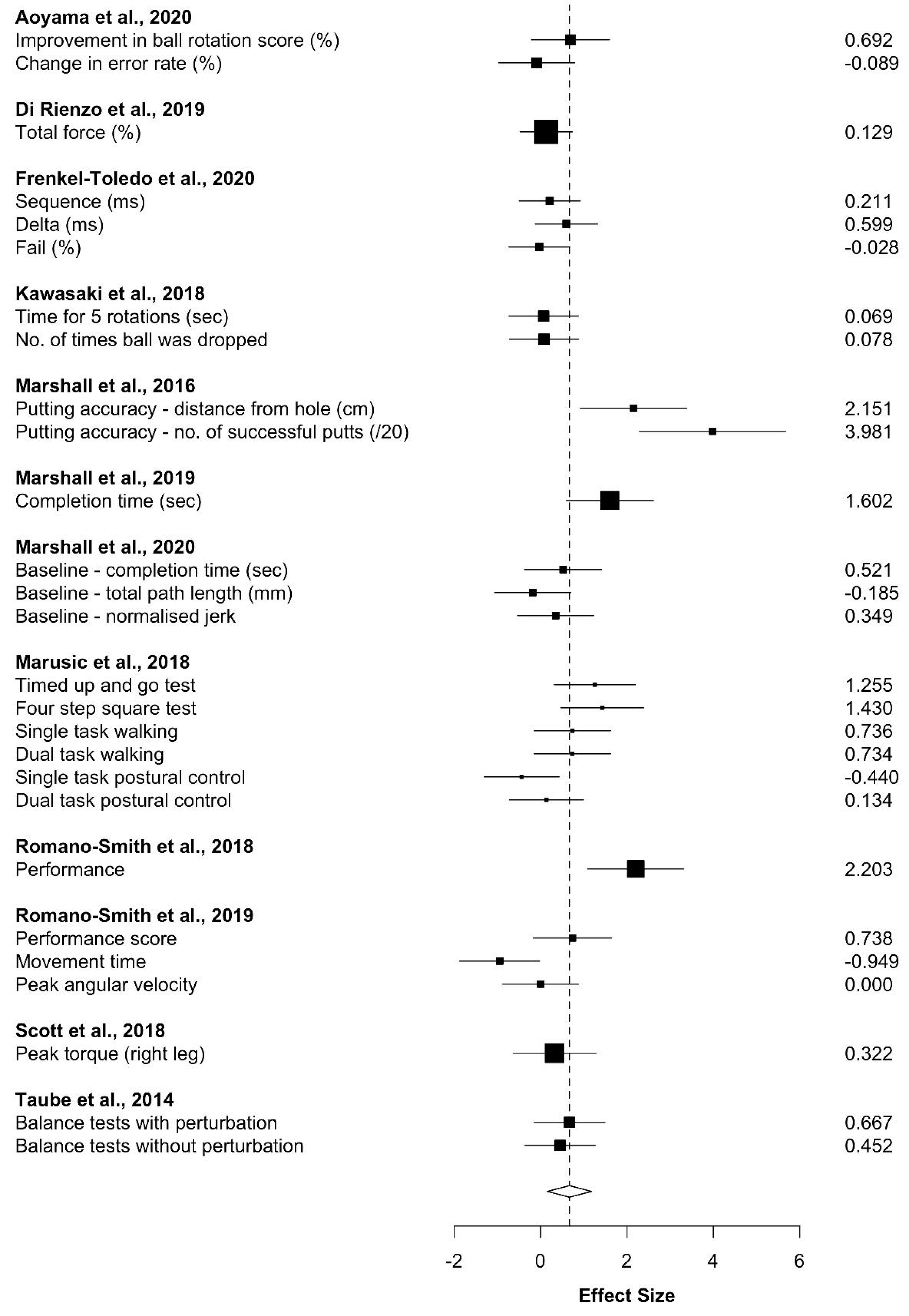
Forest plot of effect sizes (*d*) for all studies included in the AOMI vs control condition comparison for the Movement meta-analysis. The combined estimate (dashed vertical line) and 95% confidence interval (hollow diamond) indicates AOMI has a medium to large positive effect on movement outcomes compared to control conditions. The size of each black square indicates the weight of the study effect size in the combined analysis. Multiple effect sizes are reported for a study if it recorded more than one movement outcome variable.

**Figure 7.**
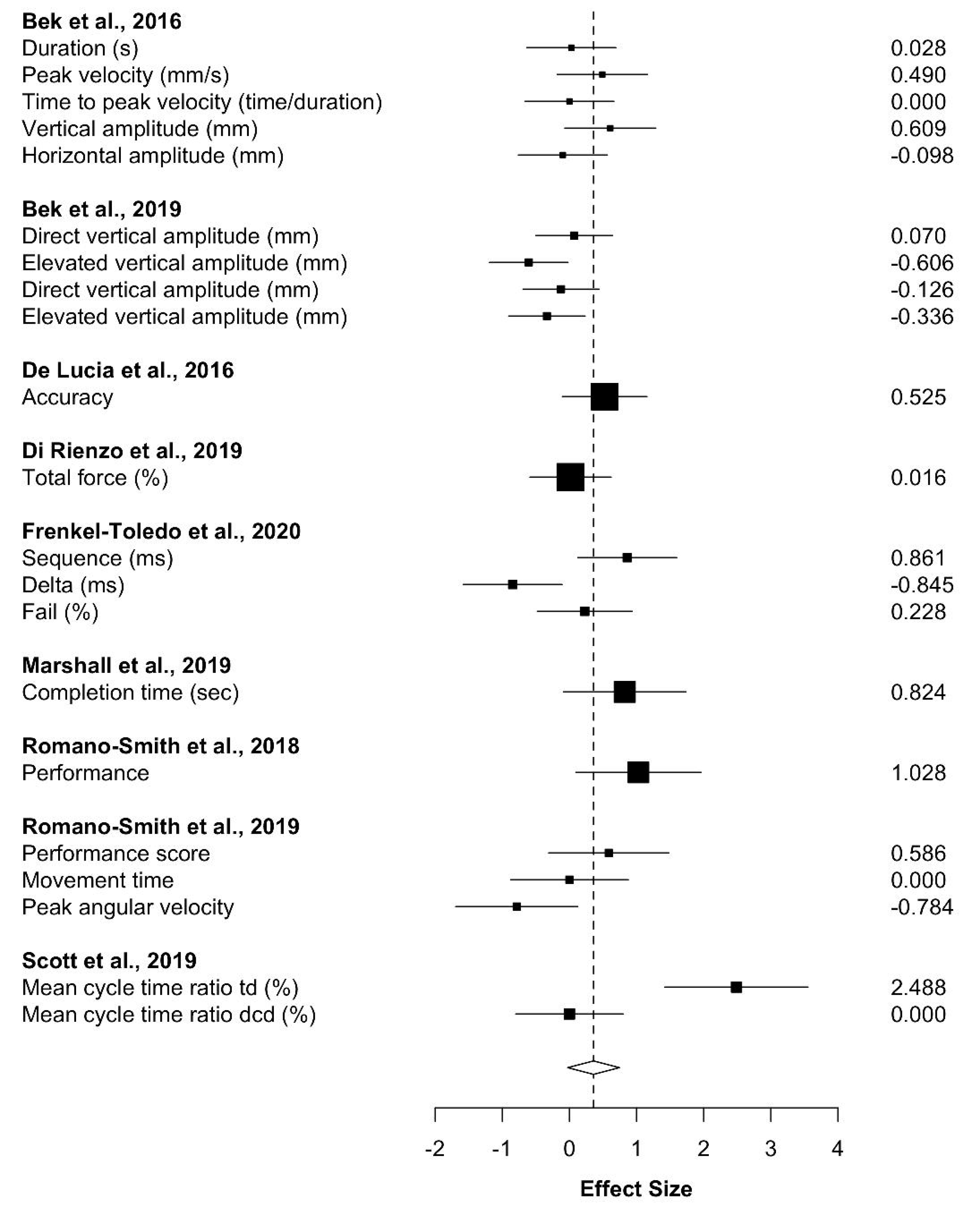
Forest plot of effect sizes (*d*) for all studies included in the AOMI vs AO condition comparison for the Movement meta-analysis. The combined estimate (dashed vertical line) and 95% confidence interval (hollow diamond) indicates AOMI has a small to medium positive effect on movement outcomes compared to AO conditions. The size of each black square indicates the weight of the study effect size in the combined analysis. Multiple effect sizes are reported for a study if it recorded more than one movement outcome variable.

**Figure 8.**
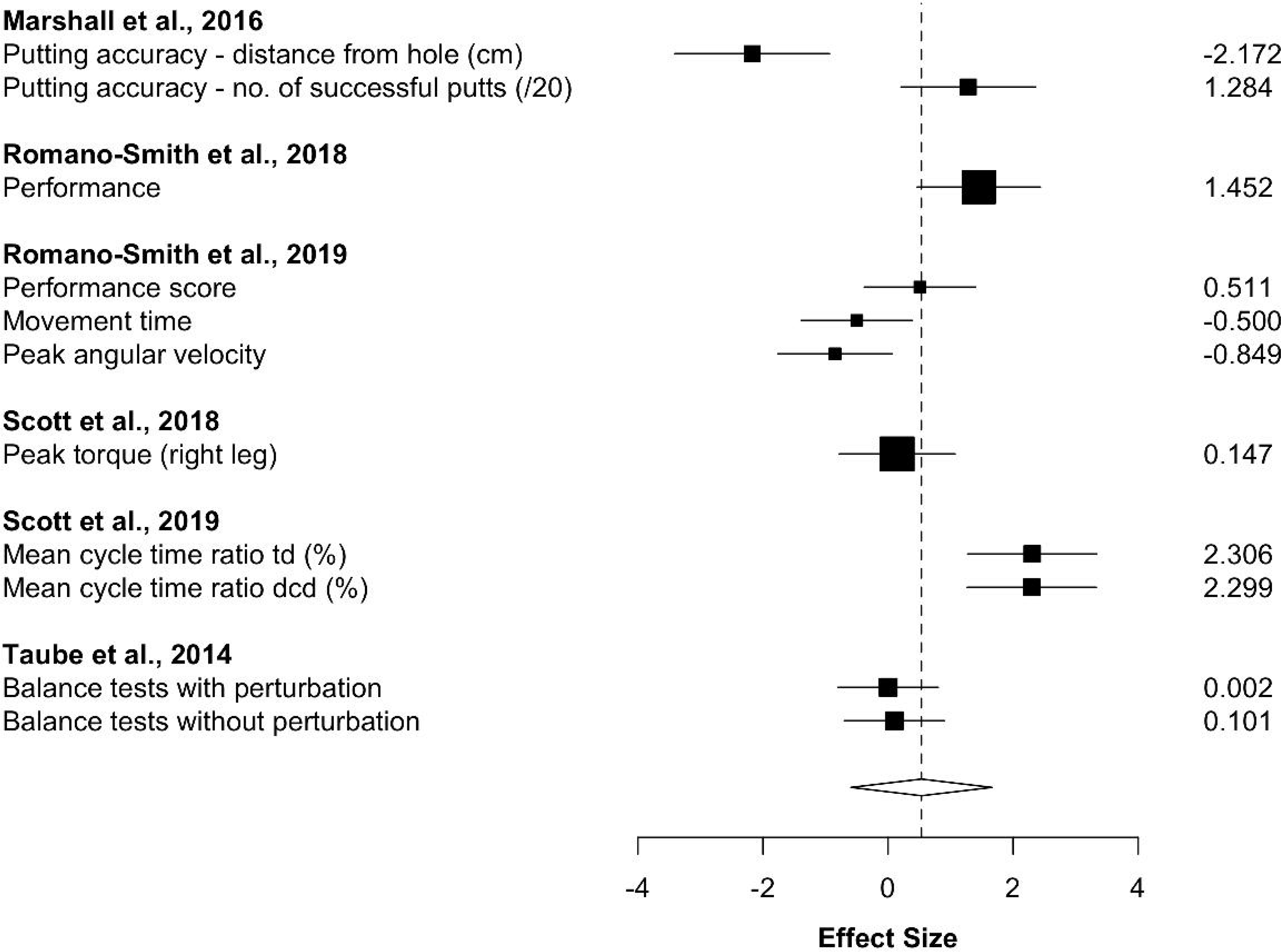
Forest plot of effect sizes (*d*) for all studies included in the AOMI vs MI condition comparison for the Movement meta-analysis. The combined estimate (dashed vertical line) and 95% confidence interval (hollow diamond) indicates AOMI has no significant effect on movement outcomes compared to MI conditions. The size of each black square indicates the weight of the study effect size in the combined analysis. Multiple effect sizes are reported for a study if it recorded more than one movement outcome variable.

**Figure 9.**
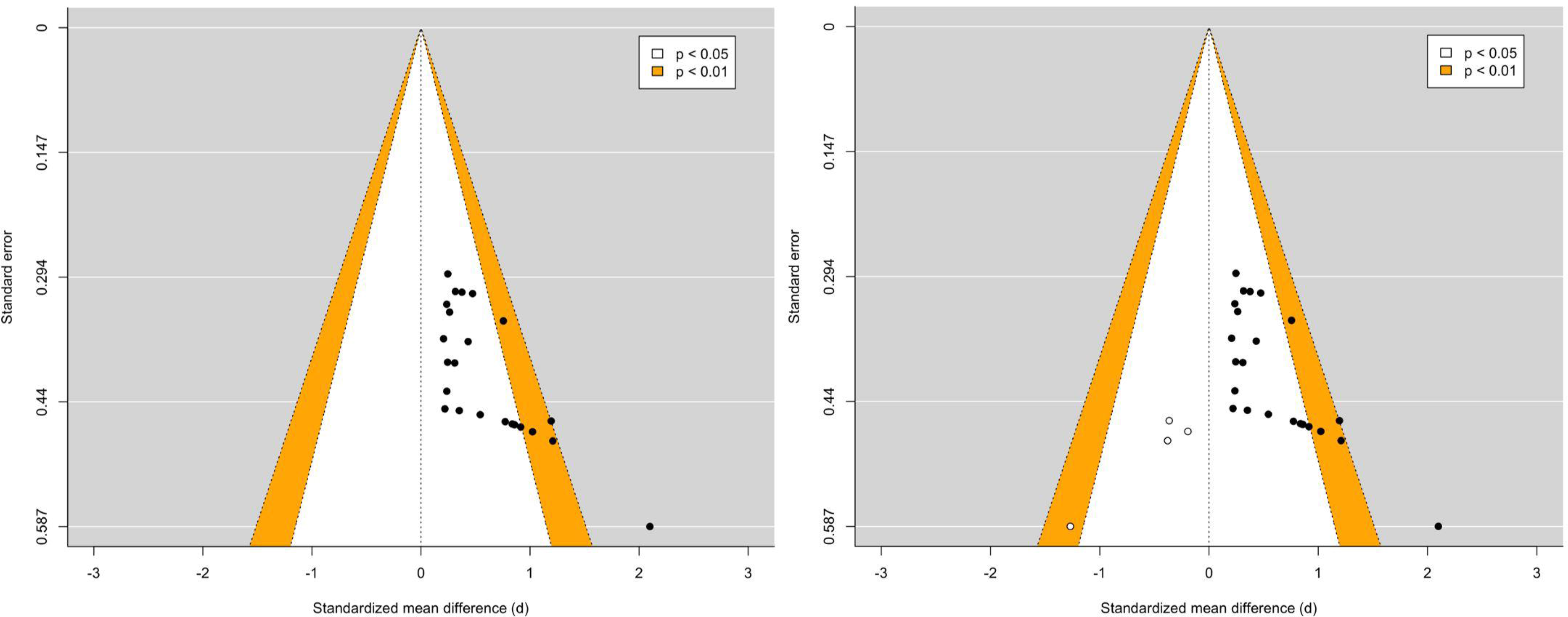
Funnel plot of effect sizes (Cohen’s *d*) versus standard error before (left) and after (right) performing Duval and Tweedie’s Trim-and-fill analysis for the AOMI vs AO comparison in the MEP meta-analysis. Black circles represent existing effects included in the MEP meta-analysis and white circles represent potential unpublished effects. The contour-enhanced funnel plots display the significance of the effects for the AOMI vs AO comparison in this meta-analysis. Individual effect sizes falling inside the white (*p* < .05) and orange (*p* < .01) funnel boundaries represent significant effects for the AOMI vs AO comparison in the MEP meta-analysis.

For the control comparison in the Movement meta-analysis, the funnel plots and trim and fill analysis showed no signs of asymmetry. Both the PET-intercept (*b0* = -0.38, *p* = 0.56) and the PET- slope (*b1* = 4.34, *p* = 0.26) were not statistically significant, suggesting that publication bias was unlikely. For the AO comparison in the Movement meta-analysis, the funnel plots and trim-and-fill analysis showed no signs of asymmetry. Both the PET-intercept (*b0* = -0.08, *p* = .88) and the PET- slope (*b1* = 2.04, *p* = .43) were not statistically significant, suggesting that publication bias was unlikely. For the MI comparison in the Movement meta-analysis, the funnel plots and trim and fill analysis showed no signs of asymmetry. Both the PET-intercept (*b0* = -0.51, *p* = .62) and the PET- slope (*b1* = 3.65, *p* = .48) were not statistically significant, suggesting that publication bias was unlikely.

## 4. Discussion

Since the early reviews introducing AOMI (e.g., Eaves, Riach et al., 2016; Vogt et al., 2013), researchers have studied both its neurophysiological and behavioral effects as a motor simulation intervention. The current paper included two meta-analyses to quantify changes in corticospinal excitability and motor skill performance for AOMI. The MEP meta-analysis collated and synthesized existing MEP amplitude data from transcranial magnetic stimulation studies as an indicator of corticospinal excitability during AOMI engagement. The Movement meta-analysis collated and synthesized existing movement outcome data from behavioral studies to assess changes in motor skill performance that result from AOMI interventions. The primary aim for both meta-analyses was to establish the effectiveness of AOMI by comparing its effects on MEP amplitudes (first meta-analysis) and movement outcomes (second meta-analysis) against those for AO, MI, and control conditions. Based on previous literature (see Eaves, Riach et al., 2016), it was hypothesized that AOMI would have a small positive effect compared to independent AO or MI, and a moderate positive effect compared to control conditions, for both outcome variables. The results of the two meta-analyses partially support this hypothesis. For the MEP meta-analysis, AOMI had a small to medium positive overall effect, a medium positive effect compared to control conditions, a small to medium positive effect compared to AO, and no significant effect compared to MI. For the movement outcome data, AOMI had a small to medium positive overall effect, a medium to large positive effect compared to control, a small to medium positive effect compared to AO, and no significant effect compared to MI conditions.

### 4.1. MEP Meta-Analysis

In the MEP meta-analysis, AOMI had a medium positive effect compared to control conditions and a small to medium positive effect compared to AO, but showed no effect compared to MI. TMS studies have consistently reported increased corticospinal facilitation for AOMI across diverse motor tasks such as simple finger movements (Bruton et al., 2020), walking (Kaneko et al., 2018), and basketball free throws (Wright, Wood et al., 2018a). From a theoretical standpoint, the finding that corticospinal excitability was facilitated for AOMI compared to control and AO but not MI conditions aligns with the propositions of the VGH that AOMI is driven by MI but may oppose the sentiments of the DASH (Eaves et al., 2012, 2014). The VGH suggests that observed and imagined actions are not co-represented, and MI is the driver for increases in motor activity during AOMI. Specifically, Meers et al. (2020) suggest the AO component acts as a visual primer, facilitating the production of more vivid images during AOMI compared to AO and MI conditions. Alternatively, the DASH proposes that concurrent representations of observed and imagined actions can be maintained as two quasi-encapsulated sensorimotor streams and that these will merge, rather than compete, when a person is overtly and covertly simulating the same action during AOMI. The merging of these two sensorimotor streams is likely to produce more widespread activity in the premotor cortex (see Filimon et al., 2015) than the AO, MI and control conditions, contributing to increased corticospinal excitability via cortico-cortical connections linking premotor and motor cortices (Fadiga et al., 2005).

The non-significant increase in MEP amplitudes for AOMI compared to MI reported in the MEP meta-analysis may be explained by the propositions of the VGH, as the increased imagery vividness for AOMI vs MI could be expected to be represented by a smaller difference in MEP amplitude between these two conditions. This difference could be expected to be negligible if those simulating actions were able to generate clear and vivid kinesthetic imagery without a visual primer, as was the case for the participants synthesized in the MEP meta-analysis (mean normalized kinesthetic imagery ability score = 67.41%, median = 70.83%, range = 55% - 76.33%). Current evidence is conflicting for the VGH and DASH accounts of AOMI (see Bruton et al., 2020; Meers et al., 2020), but both hypotheses offer feasible explanations for the impact of AOMI on the motor system and thus warrant further systematic investigation.

### 4.2. Movement Meta-Analysis

In the Movement meta-analysis, AOMI had a medium to large positive effect on movement outcomes compared to control conditions and a small to medium positive effect compared to AO conditions. Such positive effects are evidenced across most studies included in the Movement meta-analysis, with movements ranging from dart throwing (Romano-Smith et al., 2018, 2019) to whole-body balance tasks (Taube et al., 2014) in both neurotypical (e.g., Aoyama et al., 2020) and neurodivergent populations (e.g., Marshall et al., 2020). The increased motor activity during AOMI, as discussed in the previous section, is a possible neurophysiological mechanism for this effect on movement outcomes. Repeated engagement in AOMI, and thus activation of the motor system, has the potential to support repetitive Hebbian modulation of intracortical and subcortical excitatory mechanisms through synaptic plasticity, in a similar manner to physical practice (Holmes & Calmels, 2008). From a cognitive perspective, AO and MI help develop mental representations that comprise cognitive information relating to movement execution (Frank et al., 2020). When executing a motor task, a person recalls the relevant mental representation and uses this to guide their movement (Frank et al., 2020). AO and MI are proposed to contribute differently to the development of such mental representations, with AO providing sequential and timing information and MI providing sensory information related to the movement (Kim et al., 2017). It is possible combining the two forms of motor simulation during AOMI allows for the effective development of mental representations of action in the long-term memory, benefitting the physical execution of a motor task (Frank et al., 2020; Kim et al., 2017; Wright et al., 2021).

In contrast, the results of the Movement meta-analysis showed that AOMI had no significant effect on movement outcomes compared to MI conditions. Robust evidence supports the efficacy of MI as an intervention to improve motor performance across settings (e.g., MI: Guillot & Collet, 2008). This null finding aligns with the effects of AOMI on corticospinal excitability when compared to MI in the MEP meta-analysis. Specifically, AOMI did not increase corticospinal excitability or improve movement outcomes when compared to MI conditions across the two meta-analyses conducted in this paper. This provides further support for the VGH account for AOMI (Meers et al., 2020), suggesting that the imagery component drives the effects of AOMI on both the motor system of the brain and subsequent adaptations to physical movement. The sample synthesized for the MI comparison in the Movement meta-analysis were more able imagers (mean normalized kinesthetic imagery ability score = 74.78%, range = 56.67% - 96.17%) than the sample synthesized for the MI comparison in the MEP meta-analysis (mean normalized kinesthetic imagery ability score = 67.41%, range = 55.00% - 76.33%). This could suggest that individuals with high imagery ability benefit less from AOMI because the visual primer provided by AO does not improve the vividness or clarity of their MI during AOMI.

The findings of the Movement meta-analysis promote AOMI as an effective alternative intervention to AO and MI as well-established approaches, but do not indicate that combining AO and MI simultaneously (i.e., AOMI) has an additive benefit towards motor performance compared to MI. It is worthwhile noting that AOMI had a small to medium positive effect on movement outcomes compared to MI (*d* = 0.53) despite the lack of significant differences reported in the Movement meta-analysis. This is an important consideration in applied settings, such as sport and neurorehabilitation, where marginal improvements in motor performance can have practical significance (Lakens, 2013). AOMI interventions are a suitable alternative to AO and MI interventions as this combined approach can address the reported limitations of using either simulation technique in isolation. The capacity to generate and maintain mental images, termed ‘imagery ability’, is a complex cognitive process that is variable within- and between-populations (Cumming & Eaves, 2018). Individuals with low imagery ability typically find it difficult to generate and control imagined content during MI interventions, an issue that is not present for AO interventions as specific movement content can be displayed via video (Holmes & Calmels, 2008). However, the effectiveness of AO interventions is dependent on the observer’s ability to attend to the most important aspects of the motor task being performed (D’Innocenzo et al., 2016). Based on current recommendations for delivering AOMI interventions (see Wright et al., 2021), AOMI has the capacity to control the visual information displayed via AO whilst directing the individual’s attention by getting them to focus on kinesthetic aspects of the movement, subsequently reducing the complexity of MI.

### 4.3. Limitations and Future Research Recommendations

#### 4.4.1. Study Reporting

A secondary aim of this paper was to explore several methodological parameters hypothesized to have a moderating effect on the impact of AOMI interventions on MEP amplitudes or movement outcomes across the two meta-analyses. This was conducted to try and understand the influence of key methodological aspects raised in early reviews on AOMI (Eaves, Riach et al., 2016; Vogt et al., 2013) and to provide recommendations for future research and delivery of AOMI interventions. Whilst moderator analyses were run in the form of meta-regression and sub-group analyses, only the sub-group analyses included the full sets of studies for each meta-analysis (i.e., 16 studies per analysis), with missing study information meaning that meta-regression analyses included 75% of the studies on average across the two meta-analyses (mean = 12 studies, min = 6 studies for kinesthetic imagery ability in the MEP meta-analysis, max = 16 studies for sample age in the Movement meta-analysis). The issue of inadequate reporting has been raised in recent meta-analyses focusing on imagery interventions, with both noting issues related to imagery ability and assessment and reporting across studies (see e.g., Simonsmeier et al., 2021; Toth et al., 2020). A recent article has provided guidance for authors to standardise and improve the quality of reporting for action simulation studies (Moreno-Verdú et al., 2022). Alongside adhering to these useful guidelines, we also recommend that authors address the apparent biases in AOMI literature made evident by the unbalanced population sizes compared in the sub-group analyses for the two meta-analyses reported in this paper. For TMS studies, we suggest researchers employ guided attentional focus, include more diverse motor tasks (i.e., gross/continuous/serial/open skills), and test the impact of the timing of TMS delivery on the effects of AOMI on MEP amplitudes. For behavioral studies, we recommend that researchers employ guided attentional focus, adhere to PETTLEP guidelines, incorporate physical practice, recruit neurodivergent populations, and include more diverse motor tasks (i.e., gross/continuous/serial/open skills) when studying the effects of AOMI on movement outcomes.

#### 4.4.1. Neurophysiological Modality

The MEP meta-analysis included in this paper synthesized MEP amplitude data from AOMI studies using single-pulse TMS. TMS has been widely used as a neurophysiological modality with AOMI as it permits the recording of muscle-specific facilitation in corticospinal excitability, an effect that has been demonstrated robustly for AO and MI (Naish et al., 2014; Grosprêtre et al., 2016).

Whilst single-pulse TMS provides an indication of activity within the motor and premotor cortices of the brain during AOMI, EEG and fMRI can be used to provide complimentary knowledge about the roles of other cortical regions during AOMI as they can measure whole-brain activity and have high temporal and spatial accuracy, respectively (Holmes & Wright, 2017). Studies have shown that AOMI leads to activity in brain regions that would not be activated directly during the delivery of TMS to the primary motor cortex (e.g., rostral prefrontal cortex; Eaves, Behmer, et al., 2016). Activity for these brain areas is not represented within the MEP meta-analysis conducted in this paper.

Consequently, there is a need to collate and synthesize data on the precise anatomical substrates involved in AOMI using neuroscientific methods with increased spatial resolution. Hardwick et al. (2018) recently performed a large-scale activation likelihood estimation meta-analysis on fMRI data for AO, MI, and movement execution to identify distinct and shared neural regions for these three action states. This approach could be adopted to advance understanding of the neural mechanisms underpinning engagement in AOMI once additional fMRI data is available for this form of simulation.

#### 4.4.2. Study Designs

To-date, the AOMI studies synthesized in the Movement meta-analysis have almost entirely explored the short-term effects of this intervention on movement outcomes, using a between-groups comparison at one time point or adopting a pre- vs post-test study design. Whilst this approach is typical in randomized controlled trials of interventions, this does not permit accurate assessment of the long-term changes that result from AOMI engagement. The benefits of AO and MI on movement outcomes are reportedly greatest during or immediately after training, with the positive outcomes gradually disappearing in the absence of simulated practice (Stevens et al., 2003; Zhang et al., 2019). However, the performance benefits of MI training are retained beyond the intervention period (Simonsmeier et al., 2021), with repetitive engagement in MI inducing neural plasticity during recovery phases when this technique is used to acquire a skill (Ruffino et al., 2017). It remains unclear if the improvements in movement outcomes associated with AOMI are maintained after the intervention is withdrawn. Future studies should draw from the methodological approaches adopted in motor learning literature (e.g., Krakauer et al., 2019) to comprehensively examine the effectiveness of AOMI when learning and improving movement outcomes for different populations and motor tasks.

#### 4.4.3. Brain-Behavior Interactions

The MEP meta-analysis synthesized data on MEP amplitudes during AOMI engagement and the Movement meta-analysis synthesized data on changes in movement outcomes after AOMI interventions. The crossover between these effects could not be analyzed in the current paper as no studies to our knowledge have collected both measures for AOMI whilst satisfying the inclusion criteria for the two meta-analyses reported in this paper. The results of the two meta-analyses show that AOMI had a positive effect on MEP amplitudes and movement outcomes compared to control and AO conditions. Collecting neurophysiological responses, such as MEP amplitudes, alongside movement outcome measures will advance our understanding of the relationship between the changes in brain activity and motor skill performance for this intervention. Such integration of measures has shown that corticospinal excitability during MI is related to the magnitude of motor cortical adaptations after MI training (Yoxon & Welsh, 2020), and that MI and physical practice lead to different changes in brain activity and subsequent movement execution after equal training bouts (Kraeutner et al., 2020). Future research should combine neuroscientific and human movement science methods to elucidate the neural mechanisms underlying improvements in motor skill performance and learning through AOMI.

## 5. Conclusion

The two meta-analyses included in this paper synthesize the existing MEP amplitude and movement outcome data for AOMI to compare its effectiveness against AO, MI, and control conditions. The results of the MEP meta-analysis report that AOMI has a small to medium positive overall effect on MEP amplitudes, as an indicator of corticospinal excitability. When compared to different conditions, AOMI has a medium positive effect on MEP amplitudes compared to control, a small to medium positive effect compared to AO, and no significant effect compared to MI conditions. For the Movement meta-analysis, AOMI had a small to medium positive overall effect on movement outcomes. When compared to different conditions, AOMI has a medium to large positive effect on movement outcomes compared to control, a small to medium positive effect compared to AO, and no significant effect compared to MI conditions. No methodological factors moderated the effects of AOMI in either of the two meta-analyses, indicating a robust effect of AOMI for both outcome variables. However, it should be noted that inadequate reporting of methodological information as well as limited variation in the current literature on AOMI may have resulted in biased comparisons being made between moderator sub-groups and low powered assessments of relationships across the two meta-analyses. Overall, the results of both meta-analyses support the effectiveness of AOMI as an alternative intervention to AO and MI, two well established interventions, as AOMI engagement addresses the limitations of using these approaches in isolation when targeting increased activity in motor regions of the brain and improvements in motor skill performance. A more methodologically diverse approach that integrates brain and behavior is needed in future AOMI research to advance the current state of knowledge for this intervention.

